# The maternal foam plug constitutes a reservoir for the desert locust’s bacterial symbionts

**DOI:** 10.1101/2020.09.15.296319

**Authors:** Omer Lavy, Uri Gophna, Amir Ayali, Shalev Gihaz, Ayelet Fishman, Eran Gefen

## Abstract

A hallmark of the desert locust’s ancient and deserved reputation as a devastating agricultural pest is that of the long-distance, multi-generational migration of locust swarms to new habitats. The bacterial symbionts that reside within the locust gut comprise a key aspect of its biology, augmenting its immunity and having also been reported to be involved in the swarming phenomenon through the emission of attractant volatiles. However, it is still unclear whether and how these beneficial symbionts are transmitted vertically from parent to offspring. Using comparative 16S rRNA amplicon sequencing and direct experiments with engineered bacteria, we provide here evidence of the vertical transmission of locust gut bacteria. The females perform this activity by way of inoculation of the egg-pod’s foam plug, through which the larvae pass upon hatching. Furthermore, analysis of the biochemical structure of the foam revealed chitin to be its major component, along with immunity-related proteins such as lysozyme, which could be responsible for the inhibition of some bacteria in the foam while allowing other, more beneficial, strains to proliferate. Our findings reveal a potential vector for the transgenerational transmission of symbionts in locusts, which contributes to the locust swarm’s ability to invade and survive in new territories.

## Introduction

Locusts (order: Orthoptera) have affected the lives of people at least since biblical times (Old Testament: Exodus 10: 4–19), and still constitute a serious agricultural threat today (FAO 2020; Zhang et al., 2019). During locust outbreaks or plagues, these highly polyphagous insects (Chapman & Joern 1990) perform long-distance migrations, devastating agriculture in large parts of the developing world (FAO 2020; Symons & Cressman 2001; van Huis et al., 2007; Cease et al., 2015). The desert locust (*Schistocerca gregaria*) is a well-known locust species, mostly originating in the African Sahel (Symons & Cressman 2001; Cease et al., 2015; Lorenz 2009). Under the appropriate conditions, *S. gregaria* swarms develop and can potentially reach the Arabian Peninsula, the Middle East, southern Europe, and even south-west Asia (FAO 2020).

These large-scale migrations comprise several consecutive generations (Symons & Cressman 2001; Skaf et al. 1990). Each generation develops through five nymphal instars into reproductive adults ((Symons & Cressman 2001). Post-copulation, females oviposit their egg pods within the soil, enveloping the eggs in a fine sheath of foamy secretion of previously unknown composition. The same secretion is deposited as a thick foam plug above the egg pods (Fig. 1). Upon hatching the young hatchlings crawl through the foam plug to reach the soil surface and start a new locust generation (Symons & Cressman 2001; Uvarov 1997; Hägele et al., 2000).

**Figure 1:**
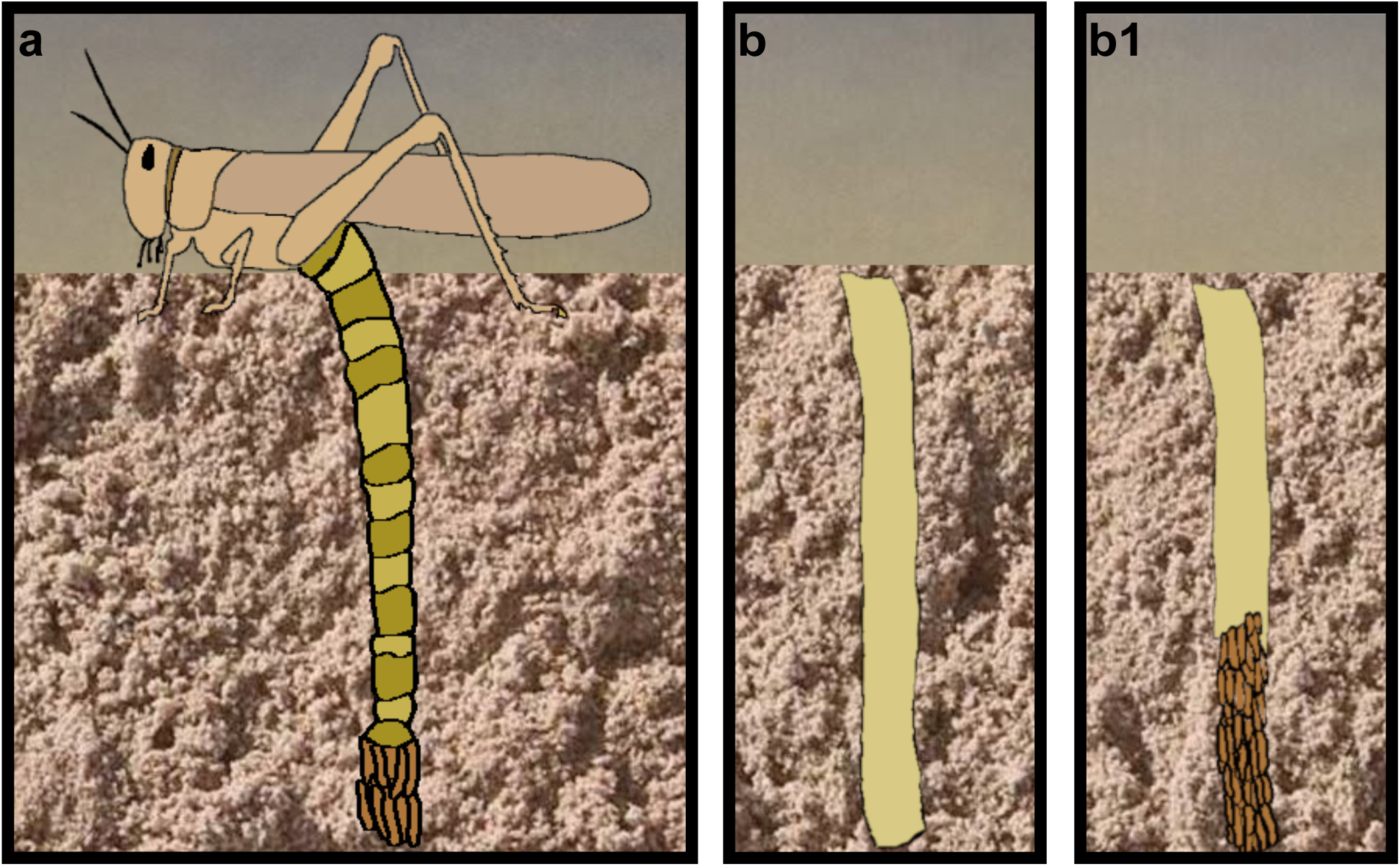
Illustrations showing **(a)** A female locust extending its abdomen while digging in order to lay eggs in the soil and laying eggs up to 14 cm deep in the soil. **(b)** The egg pod is enveloped in a secretion of foam. **(b1)** The foam sheath surrounding the eggs, partially removed to illustrate the foam plug above the eggs.

An important aspect of the desert locust biology is that of its symbiosis with the bacteria inhabiting its hindgut. These bacteria have been shown to augment the locust’s immunity through colonization resistance (Dillon & Charnley 1995, 2002; Dillon et al., 2000, 2005), and were also suggested to contribute to maintaining the swarm’s integrity through the emission of attractant volatiles (Dillon & Charnley 2002; Dillon et al., 2000, 2002). Although *S. gregaria* does not engage in an obligate interaction with any specific bacterial species, it is consistently associated with bacteria of the families *Enterobacteriaceae* and *Enterococcaceae* (Dillon & Charnley 2002, Shi et al., 2014; Lavy et al., 2019, 2020). It has been traditionally believed that the locust’s acquired bacteria are strictly environmentally-determined (Dillon & Charnley 2002; Salem et al., 2015). However, migrating *S. gregaria* in their swarming phase encounter a variety of different environnements, plants, and oviposition sites, which vary in terms of the bacteria to which the locusts are exposed (Cease et al., 2015; Uvarov 1977; Popov 1958; Pener & Simpson 2009; Soussi et al., 2015). Hence, it is unlikely that the habitat of the newly-hatched individuals can be the sole source of the bacterial agents that eventually play the pivotal physiological roles noted above. Rather, we hypothesized that the locusts possess a core microbiome that is vertically transmitted across generations; and that the effect of such a mechanism could be amplified through a selectivity trait of the locusts or through the transmission mode of the symbionts.

There are numerous examples of insects that inoculate their offspring with beneficial bacteria, through different mechanisms. The burying beetle (*Nicrophorus vespilloides*), for example, manipulates the bacterial composition of the carcasses upon which its larvae are reared (Shukla et al., 2017), and inoculates its young with advantageous gut bacteria (Wang & Rozen 2017, 2018). Other examples include brood-cell smearing with protective vertically transmitted *Streptomyces* by digger wasps (Kaltenpoth et al., 2005,2019; Koriss et al., 2010; Engl et al., 2018); and symbiont-enclosing capsules deposited by *Plataspidae* (suborder: Heteroptera) females to enable symbiont acquisition by their hatchlings (Fukatsu & Hosokawa 2002). These and other inoculation mechanisms have been thoroughly reviewed in the past (Salem et al., 2015; Onchuru et al., 2018).

We recently reported the same operational taxonomic unit (OTU), assigned to *Enterobacter* (*Enterobacteriaceae*), as being present in gregarious and solitary locusts (both laboratory-reared) across several generations (as well as in field-collected *S. gregaria*; Lavy et al., (2019)). This finding suggests transgenerational symbiont inoculation in the desert locust. Here, we examined whether locusts vertically transmit beneficial bacterial agents, and uncovered a mechanism by which bacteria may be inoculated across generations, contributing to the successful migration of locusts to new territories.

## Results

Since it had been shown previously in the desert locust that bacterial agents are not transmitted vertically through the germline (Charnley et al., 1985), we employed 16S rRNA gene amplicon sequencing to compare the bacterial composition of gregarious *S. gregaria* females to that of their offspring and to the immediate environment of their egg pod (i.e. foam and sand) (data are openly available in the SRA archive: PRJNA598984).

In an attempt to explore uncover the possible routes of symbiont transmission, our first analysis compared the bacterial composition of hatchling and pre-hatched siblings derived from the same egg pod: hatchlings (newly-hatched locusts that had climbed to the soil surface; n=16 egg pods), and pre-hatchlings (viable, healthy-looking embryos, excavated minutes before their estimated hatching time; n=11 egg pods). The individuals resulting from these two treatments have been shown to differ in both their bacterial diversity (mean values of Shannon diversity index: 2.09 and 1.28 respectively; Fig. 2) and in their overall bacterial composition (genus level, Bray-Curtis-based analysis of similarities-’ANOSIM’: R=0.15, *p*=0.024). Assuming that no bacterial transmission occurs through the germ line (Charnley et al. 1985, Dillon and Charnley 2002) these differences in bacterial composition found between the two treatments suggest that the hatchlings acquire bacteria from their immediate environment post-hatching, with either the foam or the surrounding sand as potential sources of inoculation. Moreover, the foam samples (n=15 egg pods) showed a significantly higher bacterial diversity in comparison to the sand samples (n=16 egg pods) (mean values of Shannon diversity index: 1.07 and 1.74 respectively, Fig. 2), indicating the foam plug (through which the offspring must crawl post-hatching), as a potential bacterial reservoir and source of inoculation.

**Figure 2:**
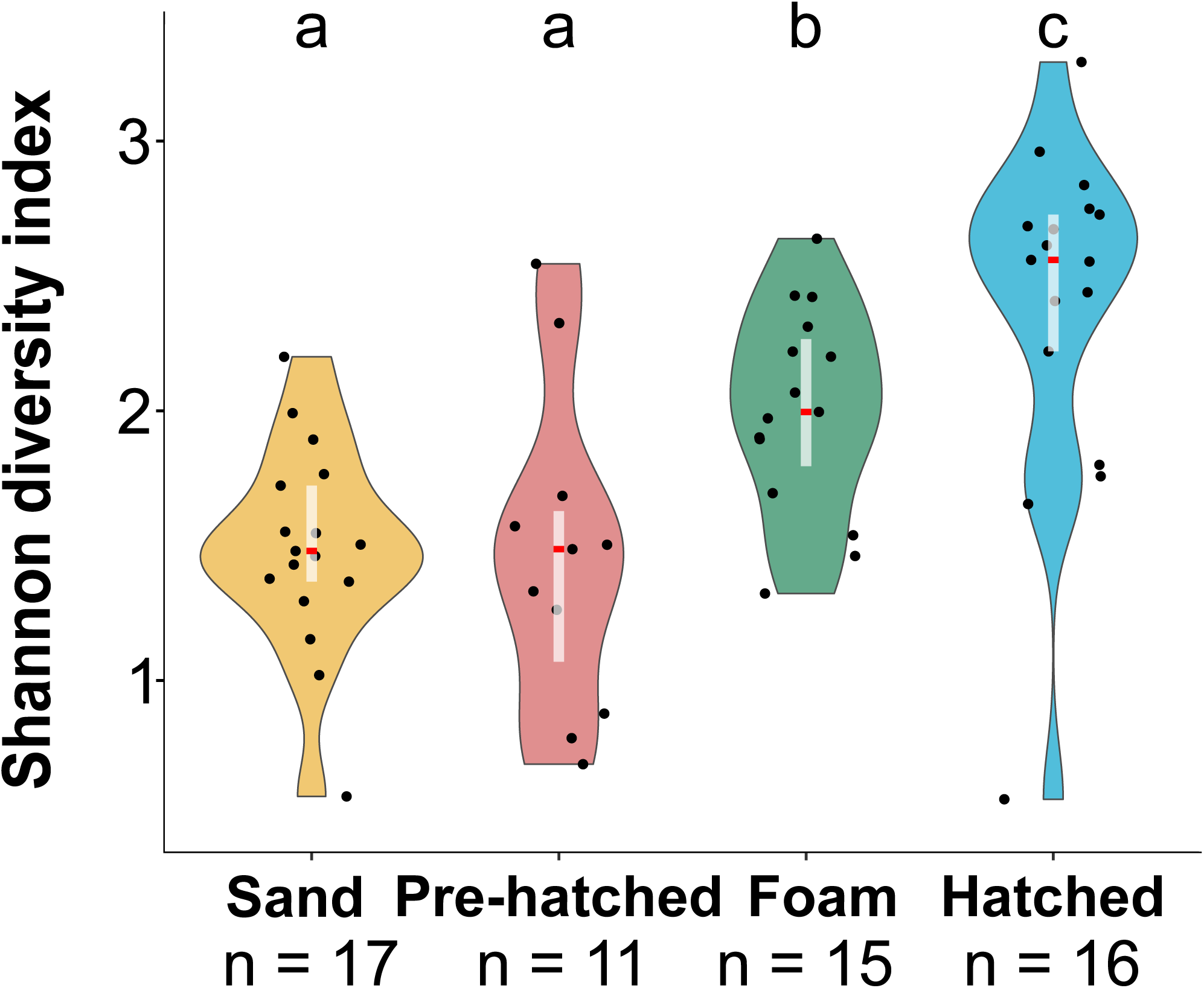
Genus-level, Shannon biodiversity indexes of the bacterial composition of the hatched and unhatched siblings, as well as the foam and sand surrounding the eggs. Differences between the groups were analyzed by Kruskal-Wallis test and Dunn’s post-hoc test. Boxes within the violin represent the range containing 50% of the data points, and the red horizontal lines denote the median.

Next, we identified 23 common amplicon sequence variants (ASVs) shared among the hindgut of the females (i.e. the mothers), the foam plug and the hatchlings, including sequences assigned to the genera *Enterobacter, Klebsiella*, and *Corynebacterium*, which have been previously shown to be associated with *S. gregaria* (Dillon et al., 2002; Lavy et al., 2019, 2020; Fig. 3b). Further analysis of the core genera (comprising the bacterial composition of at least 80% of each sample type) revealed that while *Corynebacterium* was prevalent in the foam and hatchling samples, it did not pass the threshold to be considered as core bacterium in the hindgut (did not reach >80% of the gut samples). Other core bacteria prevalent in the female hindgut, such as *Weissella*, were absent from both the foam and hatchlings (Fig 3). Nevertheless, a comparison of ASVs in full sets of mothers and their specific foam plugs and offspring, revealed the presence of the same ASV assigned to the genus *Corynebacterium* (ASV 2), as being shared among these samples in 10 out of 14 sets (∼71%, Table 1).

**Table 1:** Bacterial amplicon sequence variants (ASVs) shared among samples of locust females, the foam plug secreted by those specific females, and the hatchlings emerging through this foam (only full sets of samples are presented, n=14 sets). The bacterium Corynebacterium (bold) is shared among 10 of the 14 data sets.

| Female ID | Number of shared ASVs | Assigned genera |
| --- | --- | --- |
| 1 | non | na |
| 2 | 2 | **\*Corynebacterium**<br>*Pseudocitrobacter* |
| 3 | 3 | **Corynebacterium**<br>*Paracoccus*<br>*Brevibacterium* |
| 4 | 1 | **Corynebacterium** |
| 5 | 1 | **Corynebacterium** |
| 6 | 2 | **Corynebacterium**<br>*Stenotrophomonas* |
| 7 | 2 | **Corynebacterium**<br>*Brachybacterium* |
| 8 | 1 | *Klebsiella* |
| 9 | non | na |
| 10 | 4 | **Corynebacterium**<br>*Klebsiella* (x2)<br>*Staphylococcus* |
| 11 | 3 | **Corynebacterium**<br>*Micrococcus*<br>*Pelomonas* |
| 12 | 2 | **Corynebacterium**<br>*Klebsiella* |
| 13 | non | na |
| 14 | 4 | **Corynebacterium**<br>*Enterobacter*<br>*Brevibacterium*<br>*Enterococcus* |

\* The *Corynebacterium* assigned ASV, is identical throughout the presented data.

**Figure 3:**
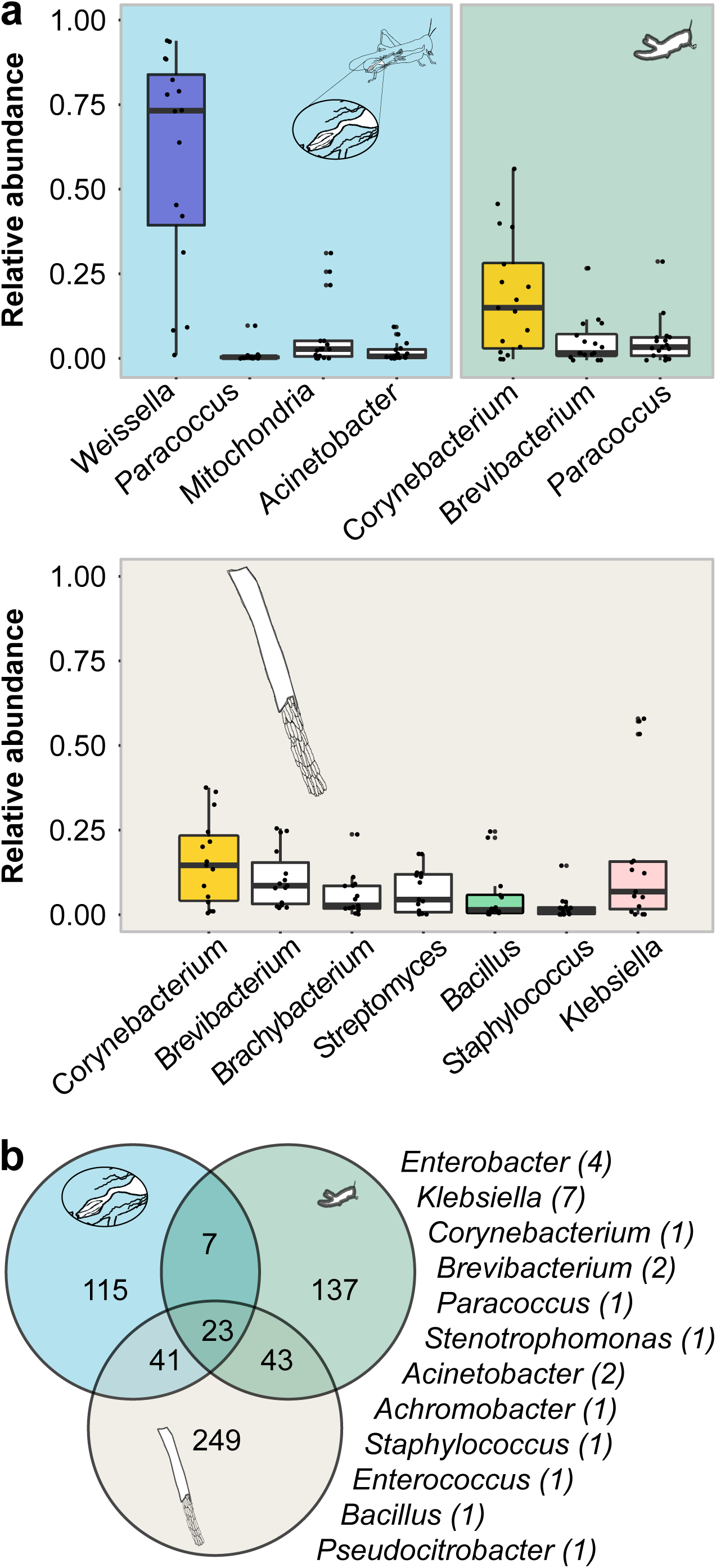
**(a)** Relative abundance of the core bacterial genera (present in minimum 80% of the samples of a specific tissue) in a female’s hindgut (left, n=16), in the hatchlings emerging through the foam (right, n=16), and in the foam (bottom, n=15). Inserts present a sketch of the sample origin (highlighted in white). Bars representing bacteria noted in the main text are colored. **(b)** Venn diagram of the dominant shared ASVs (> 75 reads per sample type) among the female’s hindgut, the hatchlings, and the foam (background color as in a, above). Taxonomical assignment of the 23 shared sequences is listed to the right of the diagram.

Further paired-sample analysis of the same data revealed a significant correlation between the relative abundance of *Corynebacterium* in specific foam plugs and the hatchlings that had crawled through these plugs (Fig. 4). Such correlation of relative abundance levels was not observed for the pre-hatching individuals and the foam plugs, nor for the hatchlings and the sand surrounding them. These findings further suggested a role for the foam plugs in the post-hatching bacterial acquisition, and that the genus *Corynebacterium* is transmitted in such a manner.

**Figure 4:**
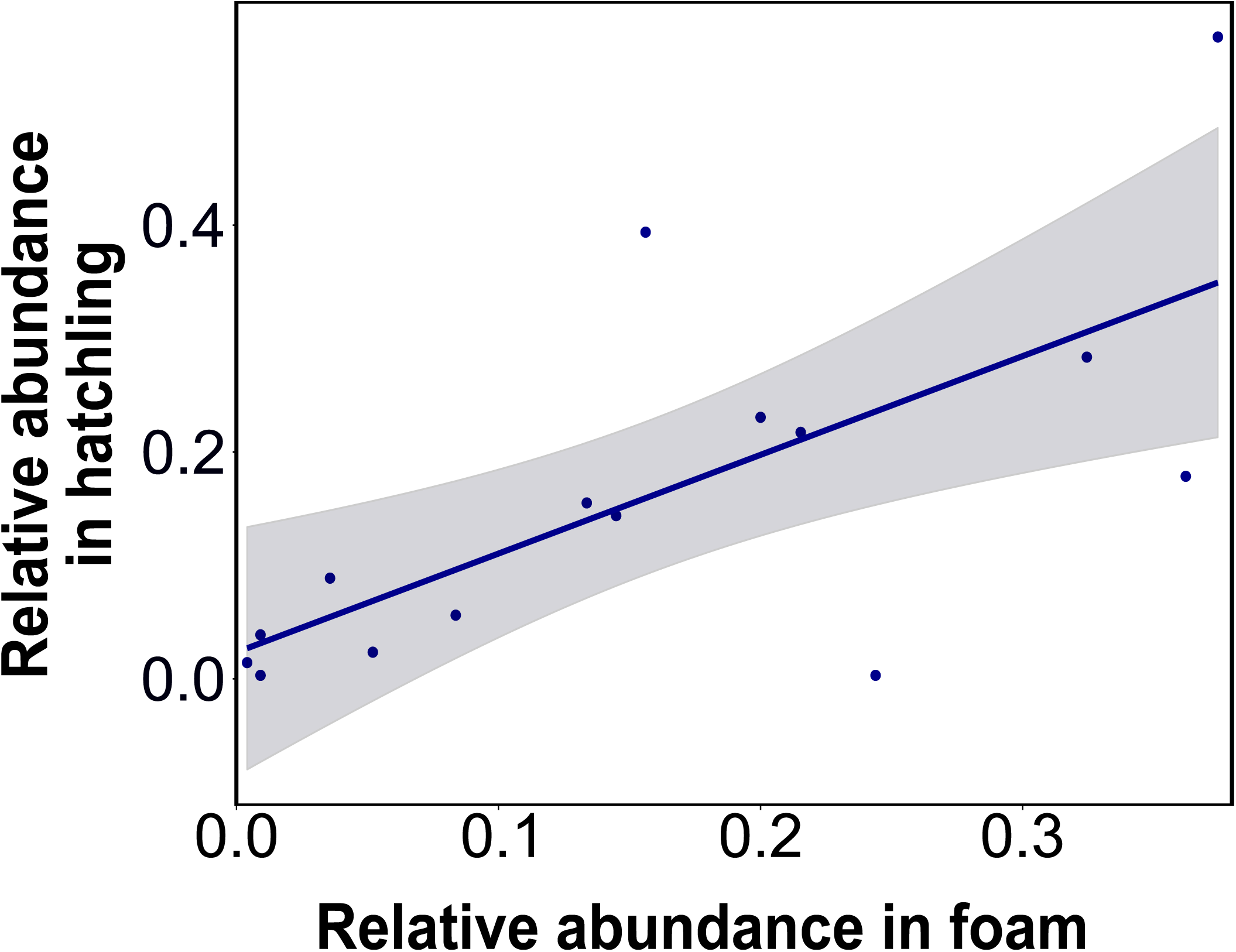
Graphical representation of the significant correlation of *Corynebacterium* relative abundance in the foam and in the hatchlings surfacing through that foam (Spearman rho: *p*=0.005, r= 0.67, n= 15). Dots represent foam and hatchlings from the same mother, and the gray area stands for the standard error.

To test the above hypothesis, we then compared the levels of *Corynebacterium* in foam-deprived hatchlings (“without foam” treatment; n=18 egg pods), which were manipulated to experience only the surrounding sand post-hatching, with that in their non-manipulated siblings (“with foam” treatment; n=19 egg pods). The latter showed significantly higher levels of *Corynebacterium* post-hatching (Fig. 5). This validated the above-noted hypothesis and also supported the hypothesized role of the foam in the vertical transmission of these bacteria.

**Figure 5:**
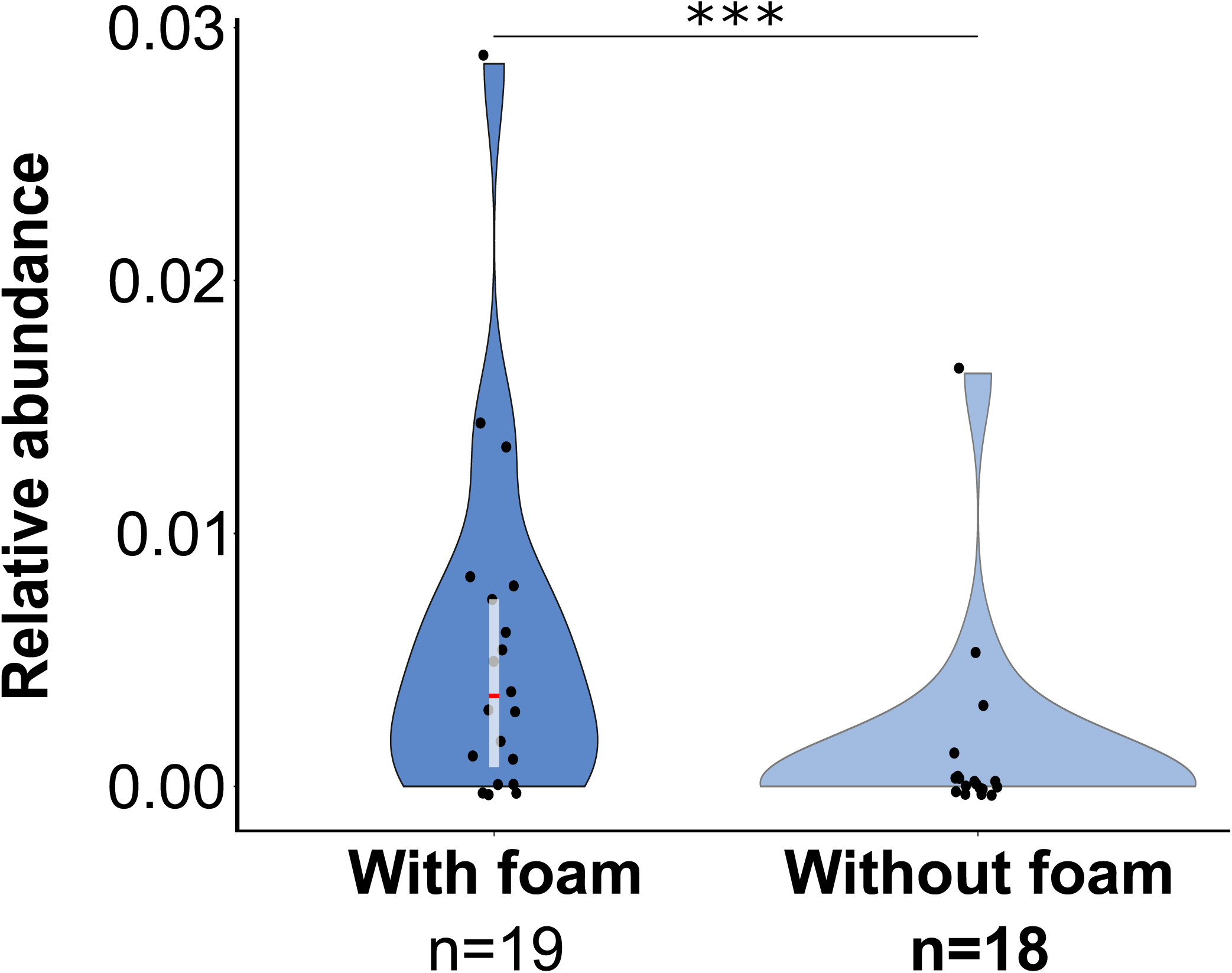
Relative abundance comparison of *Corynebacterium* in locust hatchlings surfacing in the with-foam and in the without-foam treatments. (Mann Whitney. *P*<0.001).

The next step in confirming our hypothesis would have been to demonstrate *in-vivo* the inoculation of the foam with *Corynebacterium*. However, despite substantial efforts, we were unable to isolate any locust-associated *Corynebacterium* from any of the animals. Therefore, in order to directly test the possibility of the female gut as a source of the foam bacteria, we inoculated locust females with a strain of *Klebsiella pneumoniae* isolated from locust feces. We genetically engineered that strain (see Methods) to contain two selectable markers that confer resistance to the antibiotics kanamycin and streptomycin. That resistant *Klebsiella* strain was found to be present in the inoculated female’s feces 7 days post-inoculation. Consequently, we concluded that it was stably maintained in the female gut and we then allowed these locusts to lay eggs in sterilized sand. On day 1 post-oviposition this *Klebsiella* strain was found in 11 out of 15 foam plugs secreted by these females, but in only 3 samples of the sand surrounding the egg pods. The engineered *Klebsiella* strain was still detectable in five foam plugs and two sand samples after 10 days.

### Chemical analyses

The absence of *Weissella* (which was very dominant in the female gut; Fig. 3a) in the foam, combined with the constant presence of *Corynebacterium* and other locust associated bacteria, such as *Klebsiella* (Fig. 3a) in the plugs, suggested some bacterial selectivity traits of the foam. To determine whether the foam plug might possess biochemical properties that select for particular microbes, we analyzed its composition. We determined the foam plugs’ total protein content by elemental analysis, and identified the protein repertoire using SDS-PAGE and MS analysis. Elemental analysis of the foam plug (Table. 2) indicated the presence of ∼0.63% nitrogen, which is translated to a 3-3.5% protein fraction. Nevertheless, this small protein component contained at least eight proteins that, out of the 42 proteins that could be identified by mass spectrometry, could be attributed to the locust’s immune system, such as a thaumatin-like protein, prophenoloxidase, and lysozyme (Tables 3), (Table. S3). The low protein content of the foam suggests that it mostly comprises carbohydrates. However, the extremely stable and hydrophobic nature of the foam prevented a carbohydrate profile analysis. Since the foam is insect-derived, we hypothesized that these carbohydrate molecules are composed of chitin or a chitin-like polysaccharide. Treating the foam with chitinase (a chitin degrading enzyme) caused it to lose its hydrophobicity and to change in shape and color (Fig. 6), thus confirming our hypothesis that chitin was a major component of the plug, but probably not the only component.

**Table 2:**
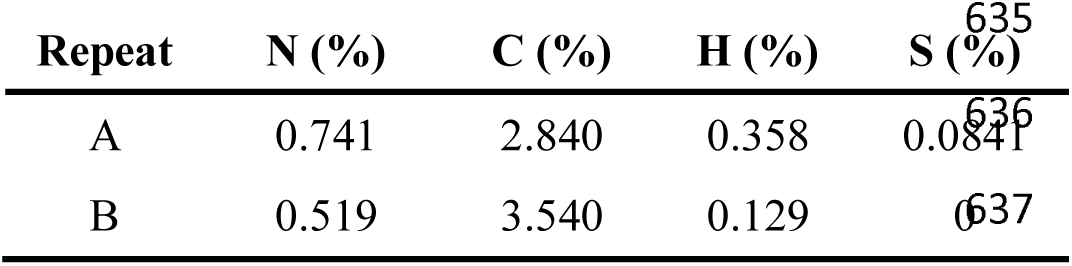
Elemental analysis of locust nest plugs.

**Table 3:**
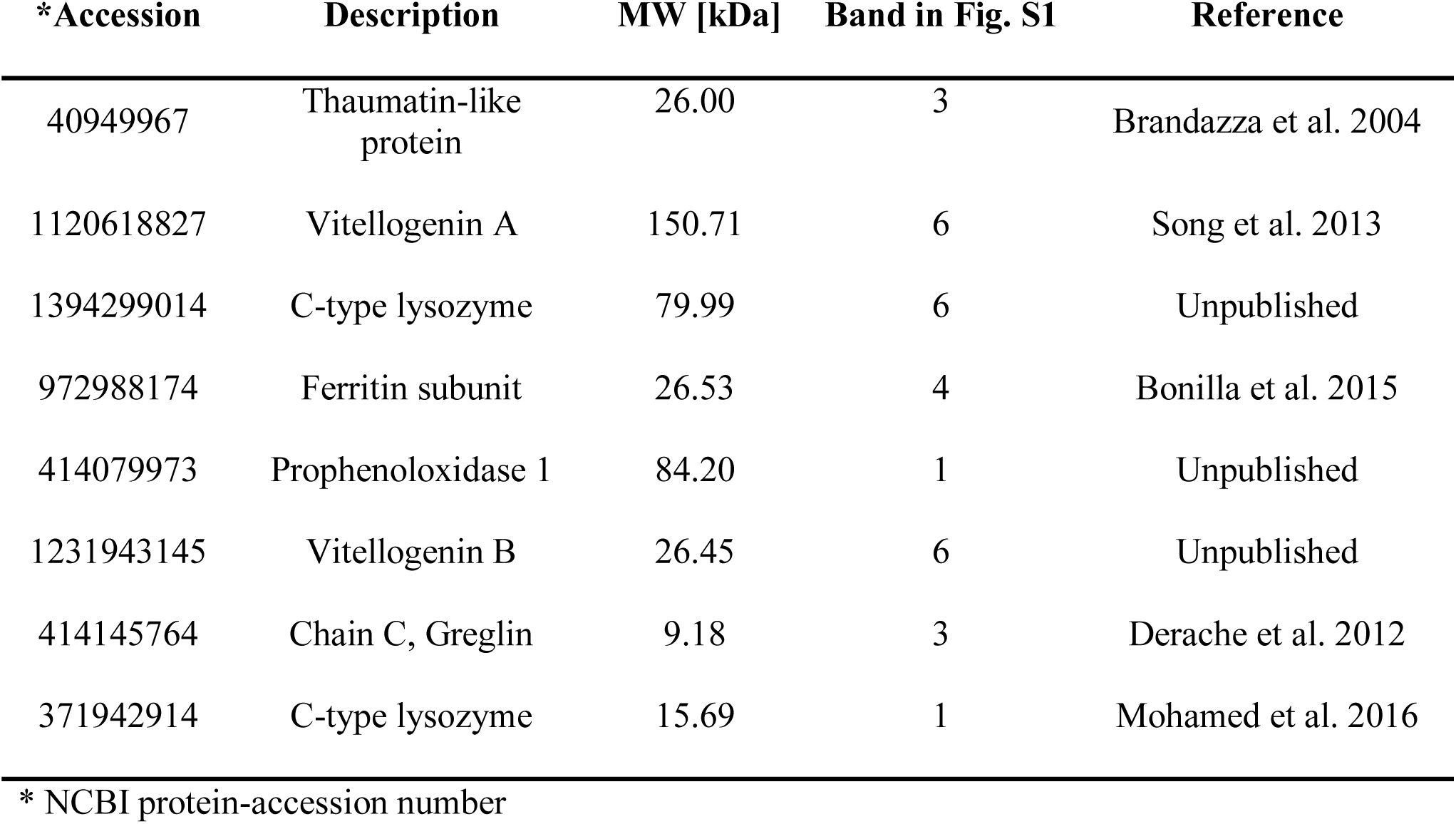
Proteins found in the foam plug and suspected to constitute immunity-conferring agents that may play a bacterial regulatory role in the foam plug. This is a partial list of 8 out of 40 proteins. Only immunity-related proteins are presented here. The full protein list can be found in Table S3.

**Figure 6:**
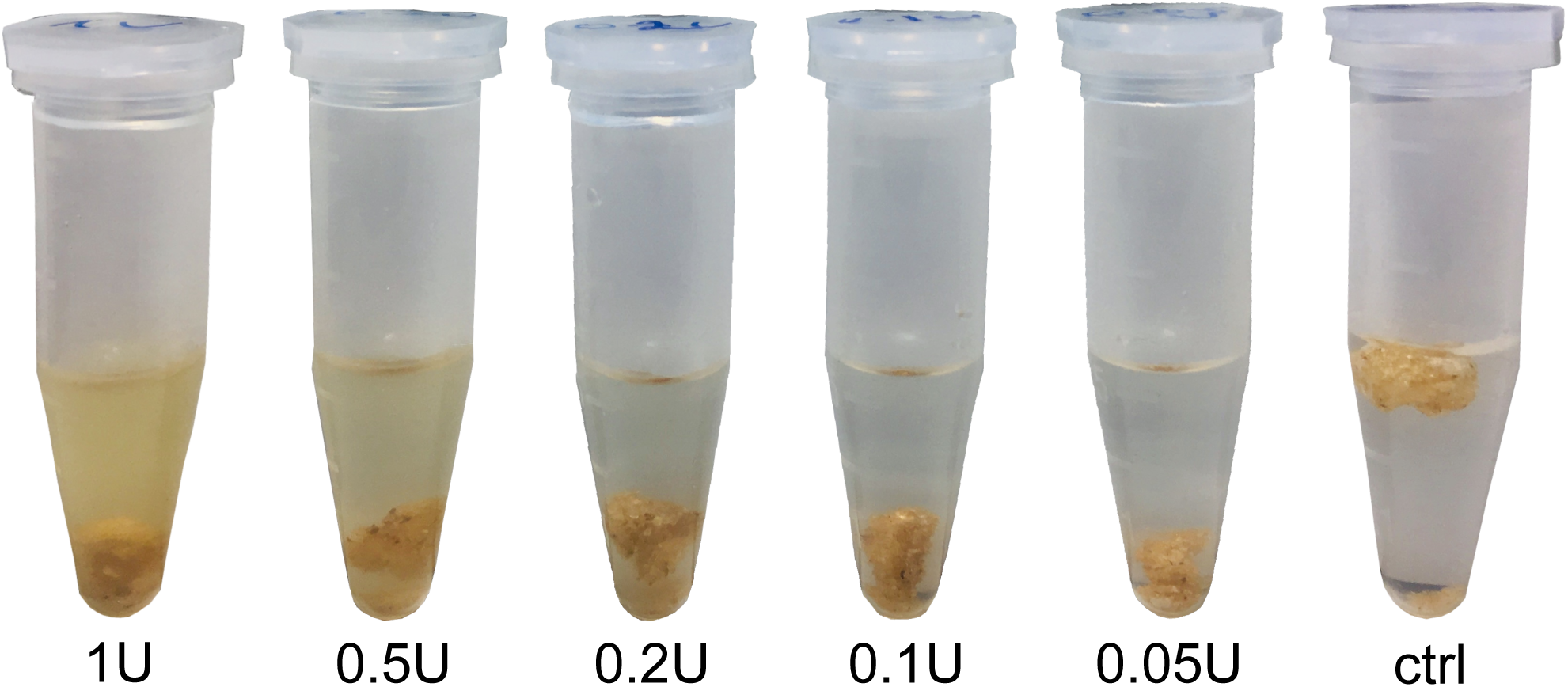
Partial hydrolysis of the foam plug mass by incubation with decreasing order of chitinase active unit (U) concentrations.

## Discussion

The success of locust swarms in covering very large distances and conquering new grounds during an upsurge (Symmons & Cressman 2001; Skaf et al., 1990) may be, at least partly, attributed to the locust’s interaction with its beneficial gut microbes, which are important for its immunity (Dillon & Chrnley 1995, 2002; Dillon et al., 2005) and may also be instrumental in maintaining locust aggregation behavior (Dillon & Chrnley 2002; Dillon et al., 2000, 2002). Though much knowledge has been acquired concerning the locusts’ bacterial symbionts (Dillon & Chrnley 2002; Shi et al., 2014; Lavy et al., 2019,2020), the mechanism that maintains these bacteria within the population and possibly transmits them from one generation to the next, has remained unclear. The commonly suggested hypothesis has been that locusts acquire their endosymbionts as hatchlings from their environment (Dillon & Charnley 2002). However, the importance of these bacteria and the diversity of feeding and oviposition habitats that females encounter while migrating (Popov 1958; Uvarov 1977; Pener & Simpson 2009; Cease et al., 2015), make it likely that they possess some sort of mechanism to ensure the inoculation of their offspring with beneficial bacteria. Here, we have shown that *S. gregaria* females deposit bacterial symbionts in the foam they secrete on top of their eggs, which in turn seems to act as a selective reservoir of these bacteria, passively inoculating the offspring post-hatching.

Our data suggest that the locust hatchlings acquire these bacterial symbionts post-hatching, while climbing to the surface through the foam plug. The bacterial ASVs, shared among the female’s gut, the foam, and the hatchlings, provide further evidence of the maternal origin of these foam-inhabiting bacteria. These findings of post-hatching inoculation are in agreement with the possibility of obtaining bacteria-free (axenic) locusts through egg surface-sterilization (Charnley et al., 1985), indicating that germ line transmission does not take place in this insect species.

However, our experiments indicate that the inoculation process is also somewhat selective: *Weissella*, for example, which is highly prevalent in the mother’s gut, was not found either in the foam or in the offspring samples at the end of the incubation period (11 days at 37°C in this case). It seems that during this period, factors in the foam enabled specific bacterial strains to proliferate, while inhibiting others. The presented data concerning the link between the *Corynebacterium* in the foam and that in the hatchlings emerging through it, further supports this hypothesis of foam selectivity; which also explains the difference in bacterial composition between the mother’s hindgut and the foam.

Although we failed to successfully cultivate locust-associated *Corynebacterium* strains, we were able to demonstrate the transmission of a genetically-modified locust-isolated *Klebsiella* strain from the female gut to the foam plug. Our results clearly indicate that locust females inoculate the foam with gut bacteria upon oviposition, offering a possible route for bacteria that originate in the female gut, to reach her offspring.

Both *Enterobacter* and *Klebsiella*, which we found to be shared among locust mothers and offspring, were also consistently found as hindgut bacterial symbionts of the desert locust in previous studies, and species of both genera were found to produce guaiacol and phenol,which are considered to be cohesion pheromones that help to maintain the integrity of the swarm (Dillon et al. 2000; Dillon & Charnley 2002). Moreover, we previously demonstrated the consistency of the same *Enterobacter* lineage through several generations of laboratory-reared locusts as well as field-collected *S. gregaria* (Lavy et al., 2019). Those findings suggested the existence of a mechanism responsible for maintaining these gut bacteria within the population, including a transgenerational inoculation mechanism, such as the foam which was the focus of the present study.

*Corynebacterium* is known to inhabit insects of different orders (e.g. Zucchi et al., 2012; Segata et al., 2016; Tobias 2016; Park et al., 2019), as well as the reproductive system of locust females (Lavy et al., 2020). Itoh et al. (1996, 1997) identified several *Corynebacterium* species that have the rare ability to utilize gaseous acetophenone as a sufficient carbon source to ensure their proliferation. Acetophenone has also been identified in the gaseous fraction emitted from foam plugs secreted by *S. gregaria* females (Rai et al., 1997). It is thus plausible that the acetophenone in the foam enriches *Corynebacterium* species that, in turn, can produce antibiotic substances (Tobias 2016; Gumiel et al., 2015). The locusts thereby “feed” the bacteria that in turn contribute to protecting the locusts’ eggs.

However, this is not the only mechanism that locusts appear to utilize to ensure the presence of specific bacteria in the foam. Mass-spectrometry analysis of the protein fraction of the foam revealed several immune-related peptides, such as lysozyme, a thaumatin-like protein, and prophenoloxidase. We speculate that the presence of these peptides in the medium surrounding the eggs provides an extra layer of protection in addition to the mechanical barrier of the foam itself. This additional protection could potentially prevent chitin-degrading pathogens from damaging the eggs, as in the case of the chironomid (*Chironomus* sp.) and *Vibrio cholerae* interaction (Broza & Halpern 2001; Laviad et al., 2016). In the locust’s case, it is possible that the gram-positive targeted lysozyme (Raglan & Criss 2017) in the foam is responsible for the absence of *Weissella* (a firmicute) in the foam and offspring, despite its dominance in the maternal gut samples. This is in contrast to *Corynebacterium* (an actinobacterium), which shows resistance to lysozyme due to its unique mycolic acids containing an outer membrane (Hirasawa et al. 2000, Toyoda et al. 2018), thus enabling it to proliferate in this environment and to be transmitted to the hatchlings as they crawl through the foam.

We have demonstrated here that chitin is a major component of the locust foam plug. The solid, hydrophobic nature of this polysaccharide probably may act as a physical protective layer against predatory organisms as well as against desiccation. To the best of our knowledge, such use of foam to protect an egg-mass, both mechanically and immunologically, has been known to date only from a few foam-nesting frog species. These frogs cover their eggs with a protein-rich foam to protect them from desiccation and pathogens (Cooper & Kennedy 2010).

In previous studies we had hypothesized that the desert locust employs some sort of mechanism in order to inoculate its offspring with beneficial bacteria; and, accordingly, that it maintains specific bacterial symbionts within the population across generations (Lavy et al., 2019, 2020). Here we have provided evidence suggesting that the foam deposited above the eggs has the potential to serve as such an inoculation mechanism. Furthermore, we provide data indicating that the foam is more than merely a reservoir for the bacteria deposited by the mothers; it is also a selective medium, potentially supporting mutualistic species while inhibiting the proliferation of others, thus maintaining a beneficial bacterial consortium for transmission across locust generations.

We are far from fully understanding the contributions of microbes to locust biology; and further *in-vivo* experiments, as well as exploration of the possible fungal and viral symbionts residing within the locusts, are needed in order to elucidate these important interactions. Nevertheless, the findings presented here provide important insights into a little addressed aspect of the locust’s ability to execute long-distance, trans-generational devastating plagues.

## Experimental procedures

To test the hypothesis of symbiont vertical transmission, we chose to combine several approaches. First, we employed 16S rRNA amplicon sequencing in order to determine the bacterial composition of locust females, their offspring, the potential bacterial inoculation vectors that constitute the eggs’ immediate surroundings (in the present study this was the sand in which the female oviposited), and the foam plug secreted onto the egg pod itself (Fig. 1c). We then used an engineered bacterium to test for *in-vivo* maternal inoculation. Finally, we applied a biochemical approach to analyze the foam’s composition, as a candidate vector for the symbionts’ maternal inoculation.

### Egg pod sampling

Gregarious locusts were reared under crowded conditions for many consecutive generations at 33±3°C and a photoperiod of 14L:10D, and fed fresh wheat grass and dry oats (for details see Lavy et al. 2019).

In order to examine the possibility of transgenerational bacterial transmission, we conducted an experiment comparing the bacterial composition of locust mothers with that of their newly-hatched and viable pre-hatched offspring.

Newly-matured females were introduced individually into11×12×14.5 cm metal cages containing two mature males and food. A 50 ml centrifuge tube (Corning, NY, United States) filled with autoclaved moistened sand was replaced daily until oviposition occurred and egg pods were detected. The females were then sacrificed and kept individually in 70% absolute ethanol, at −20°C, until tissue sampling.

The tubes containing egg pods were incubated at 37°C. Around day 11 of incubation (the typical duration of embryonic development under these conditions) the tubes were meticulously examined for the presence of hatchlings. Upon detection (within 30 min of their emergence onto the sand surface) three hatchlings were collected into a 1.5 plastic tube using sterile forceps (each pooled hatchling group is considered as one sample). The egg pod was then excavated and three healthy-appearing locust-bearing eggs were collected in the same manner.

The pooled hatchlings and the unhatched larvae were sacrificed, washed, and vortexed five times × 1 minute in filtered saline (0.9% NaCl) to remove unattached external bacteria (Fukatsu & Hosokawa 2002), and stored in 70% absolute ethanol at −20°C until further use. Samples of the foam and the surrounding sand were also collected from each tube containing an egg pod and stored in the same manner, allowing us to assign hatched larvae to their unhatched siblings and to the foam and the sand from the same tube (i.e. the same egg pod).

### Controlling for the foam as an inoculation source-with and without foam treatment

Sexually-mature females were housed individually and supplied with sand-filled oviposition tubes as described above. Prior to autoclaving, the sand tubes were cut longitudinally and then re-attached. After the female had laid eggs, the egg pods were incubated at 37°C for five days. On day 5 post-oviposition, the tubes were opened along the pre-made cut and some of the eggs were removed and placed in a fresh similar tube of autoclaved sand. The original egg-containing tubes were resealed and incubated. The eggs that had remained in the original tube gave rise to hatchlings that surfaced through the foam plug (foam treatment), while the hatchlings emerging from the relocated eggs surfaced only through a layer of moistened sand (without foam treatment). Upon surfacing, the hatchlings were collected in groups of three, washed five times as described above, and stored in 70% absolute ethanol at −20°C until further use.

### Gut sampling of females

The locusts’ wings and limbs were dissected out and their body surface was sterilized by submerging in 1% NaOCl solution for 2 min followed by two consecutive washings in fresh double-distilled water. The bodies were then dissected aseptically (according to the protocol detailed in Lavy et al. (2019)) and their hind-gut collected. The excised samples were kept individually in 70% absolute ethanol at −20°C until DNA extraction.

### DNA extraction and sequencing

Ethanol was removed and bacterial genomic DNA was extracted using the “Powersoil” DNA isolation Kit (Mo Bio Laboratories Inc., Carlsbad CA, United States), according to the manufacturer’s instructions, using 60 µl for final DNA elution. To determine bacterial composition, polymerase chain reaction (PCR) of hypervariable areas V3 and V4 of the prokaryotic 16S rRNA gene was performed on the extracted DNA; using a universal primer containing 5-end common sequences (CS1-341F 5’-ACACTGACGACATGGTTCTACANNNNCCTACGGGAGGC AGCAG and CS2-806R 5’-TACGGTAGCAGAGACTTGG TCTGGACTACHVGGGTW TCTAAT). PCR conditions: initial step of 94°C for 2 min, followed by 30 PCR cycles of denaturation at 94°C for 30 sec, annealing at 50°C for 30 sec and extension at 72°C for 30 sec, ending the reaction with 4 min at 72°C. The reactions were performed using the PCR master mix Go Taq^®^ Green Master Mix (Promega Corporation, Madison, WI, United States). PCR product validation was conducted by agarose gel 1% electrophoresis, and validated samples were sent for deep sequencing of the amplified amplicons (∼430 bp per read), conducted on an Illumina MiSeq platform at the Chicago Sequencing Center of the University of Illinois. Tubes containing all reagents but lacking samples, were added to every sequencing batch to rule out contamination.

### Data analyses

Demultiplexed raw sequences were quality filtered (bases with a PHRED score < 20 were removed) and merged using PEAR (Zhang et al., 2014). Sequences of less than 380 bp (after merging and trimming) were discarded. Data were then analyzed using the Quantitative Insights Into Microbial Ecology (QIIME) package (Caporaso et al., 2010) and Vsearch (Rognes et al., 2016) was used for chimera detection and elimination. Merged and trimmed data were additionally analyzed with the DADA2 pipeline (Callahan et al., 2016) to infer exact sequences for amplicon sequence variant (ASV) analyses. Chloroplast and *E-coli* sequences were excluded from the downstream analysis due to the known gut content, and to our inability to completely avoid Master Mix derived *E-coli* fragments. Average read depth was 21,910 seqs/sample (range 1-56,830). Data were rarified to 980 seqs/sample prior to analysis. The “with foam” and “without foam” sections were analyzed separately, and the data of this part were rarefied to 1100 seqs/sample. All statistical analyses were conducted using R v.3.4.1. (R core team 2013). Bray-Curtis based Analysis of similarities-Anosim, principal coordinate analysis (PCoA), and Spearman’s rank correlations were carried out using the vegan 2.4-3 package (Oksanen et al., 2008).

### In-vivo foam inoculation

We used a strain of *Klebsiella pneumoniae* isolated from locust females and introduced it with two different antibiotic resistance markers to test for *in-vivo* bacterial transmission from the female gut to the egg pod and its surroundings, according to the following steps:

#### 1. Electroporation of plasmid for kanamycin resistance

Locust-isolated *K. pneumoniae* (top hit type-strain: ATCC 13884(T), similarity: 99.76%), grown overnight, was inoculated into a fresh lysogeny broth (LB) and incubated at 37°C for 90 min. The cells were then chilled on ice, harvested by two rounds of centrifugation (4000 rpm for 10 min) and washed with double-distilled water, followed by resuspension in 10% glycerol. 60 µl of cell suspension was mixed with 2 µl of pUA66 (a non-conjugative, low-copy plasmid, Zaslaver et al., 2004) and electroporated. The cells were then transferred into 1 ml LB broth and incubated at 37°C for 1 h, followed by plating on a selective substrate (final concentration of kanamycin: 50 µg/ L) to select for the resistant phenotype of pUA66.

#### 2. Streptomycin resistant mutation induction

1 ml of pUA66 harboring *K. pneumoniae* LB culture was centrifuged, washed, and resuspended in saline 0.9% NaCl. 100 µl was then applied onto LB plates containing both kanamycin (50 µg/ml) and streptomycin. (100 µg/ml). A *K. pneumoniae* colony that had been formed during overnight incubation at 37°C was applied onto a fresh selective plate to maintain the resistant strain.

#### 3. Resistance stability

The engineered bacteria were cultured in antibiotic-free LB at 37°C. Every 24 hours 100 µl were transferred to a fresh 2 ml LB medium and an additional 100 µl were applied onto selective plates overnight. Suspected *Klebsiella* colonies from each plate were boiled for 10 min in 20 µl of double-distilled water. 1 µl of the boiled mixture was then used for diagnostic PCR with specific pUA66 primers (forward primer, 5’-CATAAGATGAGCCCAAG-3; reverse primer, 5’-GTCAGTACATTCCCAAGG-3) to verify pUA66 presence. PCR conditions were: initial step of 94°C for 2 min, followed by thirty PCR cycles of denaturation at 94°C for 30s, annealing at 50°C for 30s and extension at 72°C for 30s. Ending the reaction with 4 min at 72°C. The enzyme used in this reaction was PCR master mix KAPA2G Fast™ (KAPA Biosystems, Wilmington, MA, United States).

#### 4. Inoculation

Fifteen mature female locusts were force-fed with 50 µl saline containing ∼ 615 × 10^6^ *K. pneumoniae* cells and transferred together to a female-only cage. One-week post-inoculation fecal pellets were collected from each individual female, applied onto a selective MacConkey agar, incubated for 24 h at 37° C, and diagnostic PCR for pUA66 presence was conducted on *Klebsiella*-suspected colonies from each plate to confirm the presence of the engineered bacteria in the females’ gut.

The females were then transferred individually to 11×12×14.5 cm metal cages containing two mature males, and fresh tubes of autoclaved sand and food were replaced daily. Within 24 hours of oviposition, a sample of the foam plug (n =15) and the surrounding tube-sand (n =15) was diluted and streaked on selective agar MacConkey plates, and incubated at 37°C for 24 h. pUA66 diagnostic PCR was performed on *Klebsiella*-suspected colonies from each plate.

### Foam biochemical analysis

Mature female locusts were kept individually in metal cages containing two mature males. As oviposition substrate they were provided with a 50 ml centrifuge tube containing chemically inert glass beads (diameter: 1 mm) (Paul Marienfeld GmbH & Co. KG, Lauda-Königshofe, Germany) saturated with double-distilled water.

The tubes were replaced daily and, when oviposition occurred, the egg pod was collected and its foam plug was removed and washed in double-distilled water to remove the glass beads. The foam was then left for 24 hours to dry at 4°C and the dry foam was kept at room temperature until chemical analysis.

For elemental analysis measurements, the foam plugs were crushed manually, the glass beads were removed, and the crushed matrix was soaked in 0.5M NaOH in 25°C until complete decomposition (clear yellowish solution). The soluble hydrolysis products were filtered from the beads, lyophilized, and analyzed. The nitrogen content was converted to protein according to the Nitrogen-to-Protein conversion factor range for insects reported by Janssen et al. (2017).

In order to analyze the foam protein content, 1.5g of crushed foam plugs (with the glass beads) were mixed with 8 ml ethanol and sealed in a 14 ml glass vial. The mixture was incubated at 60°C for 96 h until the foam had almost completely decomposed. Ethanol evaporation was performed in an 80°C dry bath and the pellet was dissolved in reducing sample buffer (containing β-mercaptoethnol) for SDS-PAGE analysis. The sample was boiled at 100°C for 10 min. Finally, SDS-PAGE analysis was performed with 15% acrylamide gel at 90V for 4 h. The six main identified bands (Fig. S1 in supporting information) were analyzed by Mass-Spectrometry (MS) in the Smoler Protein Research Center at the Technion, Haifa. The screening of protein results was performed against the *Acrididae* and *Locusta* protein data-bases. Only identified peptides that passed the False Discovery Rate (FDR) correction with a 99% confidence interval were used for further analysis.

Since the samples could not be dissolved in a manner that would enable carbohydrate analysis, we incubated 0.04 g of sand-coated foam plugs in 50 mM potassium phosphate (pH 6.0) buffer with decreasing chitinase concentrations (Sigma-Aldrich, St. Louis, MO, USA), in order to determine chitin or chitin-like presence in the foam.

## Acknowledgments

We would like to express the authors’ gratitude to Dr. Leah Reshef for her informed suggestions and her important help in data analyses.

## Author contributions

OL, AA, and EG conceived the study and designed the experiments. OL conducted the experiments and analyzed the data. UG oversaw the microbial ecology experiments and guided data analysis. AF and SG conducted the biochemical analyses. OL wrote the first draft of the manuscript, and all authors contributed substantially to the revisions.

**Figure S1:** SDS-PAGE visualization of proteins extracted from the locust foam plug. Two lanes represent duplicates and red squares represent the band numbering used for MS analysis.

**Figure S2**: Relative abundance of the core bacterial genera (minimum of 80% presence in the tissue samples) of the pre-hatching samples (n=11); and the sand samples (n=16). Bacteria noted in the main text are uniformly colored.

**Table S3:** Full protein profile of the foam plug.

