## Supplementary figures and images for "The maternal foam plug constitutes a reservoir for the desert locust’s bacterial symbionts"

### Figure S1

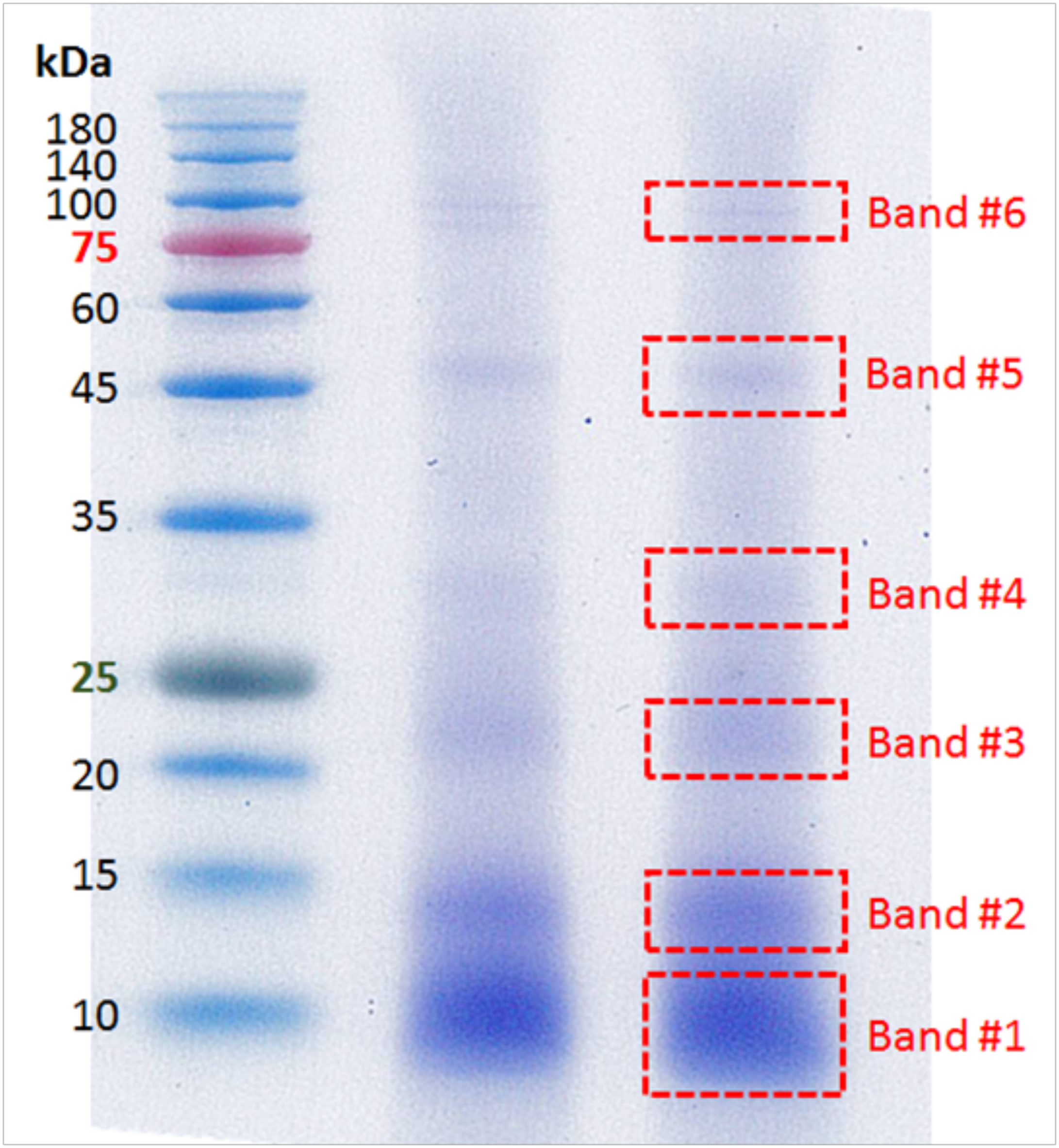

### Figure S2

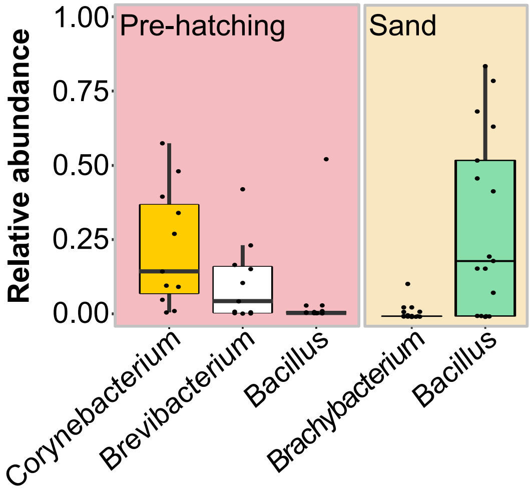
