## Supplementary material for "The maternal foam plug constitutes a reservoir for the desert locust’s bacterial symbionts": Table S3

| **Accession** | **Description** | **MW [kDa]** |
| --- | --- | --- |
| 359843282 | Actin | 41.8 |
| 40949967 | Thaumatin-like protein 1 | 26.0 |
| 1120618827 | Vitellogenin a | 150.7 |
| 1394299014 | C-type lysozyme | 80.0 |
| 972988174 | Ferritin subunit | 26.5 |
| 414079973 | Pro-phenoloxidase 1 | 84.2 |
| 1231943145 | Vitellogenin b, partial | 26.5 |
| 264667510 | Putative histone h4, partial | 6.6 |
| 1227110482 | Histone h2b | 13.6 |
| 40365371 | Angiotensin converting enzyme, partial | 72.8 |
| 146196714 | Apolipophorin precursor | 371.6 |
| 414145764 | Chain c, greglin | 9.2 |
| 972988188 | Imaginal disc growth factor 4, partial | 25.6 |
| 18874389 | Elongation factor-1 alpha | 50.4 |
| 972988176 | Membrane metallo-endopeptidase-like protein | 72.4 |
| 760244892 | Histone 3, partial | 9.2 |
| 972988208 | Angiotensin-converting enzyme | 74.7 |
| 514833443 | Mitochondrial f1-atp synthase alpha subunit | 59.5 |
| 972988140 | Midline fasciclin | 92.1 |
| 1142095214 | Mitochondrial f0f1-atp synthase subunit beta | 56.1 |
| 371942914 | C-type lysozyme | 15.7 |
| 972988198 | Lipase 3 | 41.7 |
| 1041581674 | Glyceraldehyde-3-phosphate dehydrogenase, partial | 2.8 |
| 157829838 | Chain a, molecular structure of an apolipoprotein | 17.2 |
| 227691 | Tropomyosin | 32.4 |
| 27802643 | Hsp70 family member, partial | 71.4 |
| 256368122 | Hexamerin-like protein 4 | 79.1 |
| 359843270 | Proteasome zeta subunit, partial | 12.4 |
| 359843274 | Rho gdp dissociation inhibitor | 23.6 |
| 443419060 | Hexamerin, partial | 23.1 |
| 586831203 | Carboxylesterase | 59.2 |
| 972988148 | Nucleoside diphosphate kinase | 18.9 |
| 972988184 | Proactivator polypeptide | 95.9 |
| 1034701705 | Sulfotransferase | 48.3 |
| 1041556027 | Slit | 164.9 |
| 1269806088 | Odorant binding protein 6 | 15.2 |
| 1269806091 | Odorant binding protein 7 | 24.3 |
| 77415606 | Hypothetical protein, partial | 13.8 |
| 605051977 | Chymotrypsin 12 | 28.4 |
| 46395577 | Glycine-proline-rich protein | 3.2 |
